# A Workflow for Protein Structure Determination from Thin Crystal Lamella by Micro-Electron Diffraction

**DOI:** 10.1101/2020.04.30.061895

**Authors:** Emma V. Beale, David G. Waterman, Corey Hecksel, Jason van Rooyen, James B. Gilchrist, James M. Parkhurst, Felix de Haas, Bart Buijsse, Gwyndaf Evans, Peijun Zhang

**Author notes:** Equal contributions (Emma V. Beale, David G. Waterman, Corey Hecksel). Correspondence: David Waterman, Gwyndaf Evans, and Peijun Zhang.

## Abstract

Micro-Electron Diffraction (MicroED) has recently emerged as a powerful method for the analysis of biological structures at atomic resolution. This technique has been largely limited to protein nanocrystals which grow either as needles or plates measuring only a few hundred nanometres in thickness. Furthermore, traditional microED data processing uses established X-ray crystallography software that is not optimised for handling compound effects that are unique to electron diffraction data. Here, we present an integrated workflow for microED, from sample preparation by cryo-focused ion beam milling, through data collection with a standard Ceta-D detector, to data processing using the DIALS software suite, thus enabling routine atomic structure determination of protein crystals of any size and shape using microED. We demonstrate the effectiveness of the workflow by determining the structure of proteinase K to 2.0 Å resolution and show the advantage of using protein crystal lamellae over nanocrystals.

## Introduction

Electron diffraction has become a powerful method for structural biologists and complements now well-established X-ray crystallography methods, such as rotation and serial data collection using synchrotrons and serial femtosecond crystallography with X-Ray Free-Electron Laser (XFEL). High quality data can be recorded from a very small number of sub-micron sized crystals using electrons. This arises from the strong interaction between electrons and matter, permitting protein crystals of only a few hundred nanometres to be used for structure determination (la Cruz et al., 2017; Nannenga et al., 2014), and in some cases just a single crystal (Clabbers et al., 2017). This technique therefore holds an advantage in particular cases where obtaining large crystals may be a bottleneck to structure determination. In addition, electron diffraction produces electrostatic potential maps which can offer unique information not available from their X-ray crystallography equivalent. These maps can reveal details about the charge states of atoms within the protein structure which in turn provide important insight into protein function (Liu and Gonen, 2018; Yonekura and Maki-Yonekura, 2016; Yonekura et al., 2015).

The strong interaction of electrons with matter means that the electron beam is not transmissible through samples greater than several hundred nanometres in thickness when using standard cryoEM operation voltages of 200-300 kV. In addition, multiple scattering and inelastic scattering events are also more probable as sample thickness increases (Clabbers and Abrahams, 2018). These non-kinematic scattering events affect the retrieval of structure factors from measured diffraction intensities and thus impact the quality of the final structure. To overcome this problem, we and others previously explored cryo-focused ion beam (cryoFIB) milling as a sample preparation method for electron diffraction experiments (Duyvesteyn et al., 2018; Li et al., 2018; Martynowycz et al., 2019; Zhou et al., 2019). We demonstrated that the integrity of the crystal was maintained after cryoFIB milling, and more importantly, diffraction images from a crystal lamella displayed minimal dynamical scattering (Duyvesteyn et al., 2018).

However, due to the limitation of a standard Ceta detector in our setup, the data collection strategy was not optimal and resulted in a model with poor refinement statistics, making it difficult to fully assess the potential of the lamella in the context of high-quality protein structure determination. Highly-specialised CMOS detectors, such as those produced by TVIPS, have previously been used for microED experiments (Shi et al., 2013). Hybrid pixel array detectors potentially provide a better detection method for protein crystals owing to their high dynamic range, low noise and radiation hard properties (Clabbers and Abrahams, 2018). However, these detectors are not yet commonly offered within standard high-resolution cryoEM imaging facilities. Furthermore, current microED data processing borrows software packages that were originally designed for X-ray diffraction experiments and are not optimised for handling systematic errors that are unique to electron diffraction data. To overcome these challenges and enable microED as a standard cryoEM method like single particle analysis (SPA) and cryo-electron tomography (cryoET), we established an integrated workflow for routine microED of protein crystals for implementation in any standard cryoEM imaging facility.

The workflow includes 1) cryoFIB milling to produce well-ordered crystalline lamellae (200-300 nm) from larger protein crystals; 2) a standard Ceta-D detector from Thermo Fisher for microED data collection; 3) microED data analysis using DIALS (Winter et al., 2018) which has now been optimised for electron diffraction (Clabbers et al., 2018). Using this workflow, the structure of proteinase K was determined to 2.0 Å resolution from a single protein crystal lamella. A comparison between data collected from nanocrystals and crystal lamellae suggests that higher quality structures can be obtained from proteinase K lamellae. Future automated microED data collection strategies, similar to those for SPA and cryoET, will be built upon this integrated system.

## Materials and Methods

### Crystallisation and grid preparation

Lyophilised proteinase K from *Tritirachium album* (Sigma-Aldrich, P2308) was solubilised in 25 mM Tris pH 7.5 to a final concentration of 50 mg/mL. Microcrystals and nanocrystals of proteinase K were then grown using the batch method following the protocol described by Beale *et al*., (2019). Crystal seeds were required to reproducibly grow crystals of a uniform size. The seed stock, crystallisation solution (20% (w/v) PEG 3350, 0.1 M ammonium chloride) and protein solution were combined in a 1:2:3 ratio. The crystallisation experiment was then incubated overnight with gentle agitation at 18°C. Different dilutions of the seed stock were used to generate crystals of different sizes where more concentrated seed stocks produced the smaller nanocrystals.

To produce seeds, a method based on the protocols described by Luft and DeTitta was used (Luft and DeTitta, 1999). Crystals with several hundred micrometres in size were grown using the vapour diffusion method. Crystals were grown in CrystalQuick™X plates (Greiner) at 19°C over approximately 18 hours. The drops contained 200 nL 50 mg/mL proteinase K in 25 mM Tris pH 7.5 and 200 nL reservoir solution (20% (w/v) PEG 3350 and 0.2 M ammonium chloride). Crystals from these conditions were harvested by aspirating with a pipette into a 1.5 mL Eppendorf tube containing 25 μL of reservoir solution and several small silicon beads. Crystals were crushed by three consecutive rounds of 30 s of vortexing followed by a 30 s incubation on ice. Microcrystals and nanocrystals of proteinase K were then grown using the batch method as described above.

For the nanocrystals, 3 μL of the batch crystallisation solutions were then applied to the carbon side of glow discharged R1.2/1.3 Quantifoil™ grids (Quantifoil Micro Tools, Jena, Germany). Excess liquid was removed by blotting for 12 s with a Vitrobot (Thermo Fisher) under 100% humidity at 20°C. The grids were held in the humid chamber for 30 s before plunge-freezing in liquid ethane. For the microcrystals, 2 μL of a 1 in 6 dilution of the batch crystallisation was applied to the carbon side of glow discharged R2/2 Quantifoil™ grids (Quantifoil Micro Tools, Jena, Germany). Excess liquid was removed by blotting for 4-6 s with a Vitrobot (Thermo Fisher) under 100% humidity at 20°C and immediately plunged into liquid ethane. All grids were stored under liquid nitrogen until required for cryoFIB milling or electron diffraction experiments.

### CryoFIB milling of proteinase K microcrystals

Milling of proteinase K crystals was carried out as previously described (Duyvesteyn et al., 2018; Schaffer et al., 2015; 2017) using a Scios™ DualBeam™ cryoFIB microscope (Thermo Fisher Scientific) equipped with a Quorum PP3010T cryotransfer system and cryostage and a custom-designed shuttle (Max Planck Institute of Biochemistry and Thermo Fischer Scientific). Briefly, plunge-frozen grids containing crystals of proteinase K with approximate dimensions 12 × 10 × 10 μm were loaded into autogrid (Thermo Fisher Scientific) compatible with FIB-SEM applications. The grids were then coated with an organoplatinum compound using the *in situ* gas injection system (GIS) of the cryoFIB instrument. Lamellae were generated through a series of milling steps, where the current of the Ga beam was decreased in a stepwise fashion from 300 pA to 30 pA. These steps corresponded to subsequent lamella thicknesses of approximately 5 μm down to 0.2 μm, respectively. In addition to the initial organoplatinum coating, the sample was sputter coated with metallic platinum post-milling using the Quorum PP3010T system (10 mA, 3 s, argon atmosphere) to reduce beam-induced charging effects.

### Microscope set up and data collection

All data were collected using a Talos Arctica™ TEM (Thermo Fisher Scientific) with an accelerating voltage of 200 kV. Low dose parallel illumination conditions were achieved through a combination of the largest gun lens and a small spot size 11, with a condenser (C2) apertures of 20 μm or 50 μm and operating in nanoprobe mode.

A dose rate of approximately 0.04 e^−^/A^2^/s was used for the continuous rotation data collection. Data collection essentially followed the method described by Shi *et al*. (2016) with some modifications. We used a small C2 aperture instead of a selected area aperture. A custom script (Supplementary Figure 3) was used to control the rotation speed and direction of the stage such that it was continuously rotating during data collection. Shortly after this work, the microED package EPU-D (Thermo Fisher Scientific) has become commercially available for automated data collection. Data were collected on a Ceta-D detector (Thermo Fisher Scientific) operating in rolling-shutter mode where each frame encompassed 0.51° of data. Data were recorded using Tecnai Imaging and Analysis (TIA) software as an acquisition series in the TIA/EMISPEC series file format (.ser).

During the initial frames of the continuous-rotation data collection, the diffraction pattern often appears blurry, probably due to the beam-induced specimen charging. This can be largely mitigated by sputter coating the sample with metallic Pt after cryoFIB milling (Schaffer et al., 2017) and/or refocusing the diffraction spots during the initial frames of continuous data collection. These initial frames were excluded from further analysis.

### Electron diffraction data processing

Following the scheme presented in Clabbers *et al*. (2018), the *dxtbx* library (Parkhurst et al., 2014) was extended to directly read the .ser format without further conversion. As expected, the calibrated camera length was affected by the diffraction lens adjustment used to focus the pattern. Given that the cell dimensions of the proteinase K crystals were known, we used these constraints to refine the camera length during data processing, which was carried out in DIALS following the method described in Clabbers *et al*. (2018).

The default background modelling algorithm in DIALS performs outlier handling based on the assumption that the observed counts in each pixel are approximately Poisson distributed. This model is not appropriate for the Ceta-D, which is not a counting detector. Indeed, some datasets suffered from a negative bias (Supplementary Figure 1). We therefore selected the ‘simple’ model for the background from the program options, which assumes a normal distribution of background counts. Integration was performed using DIALS 1.10, with an improved profile-fitting algorithm that is robust in the presence of negative-valued pixels.

The data were cut to a resolution where CC_1/2_ in the outer resolution shell was equal to or higher than 0.5. It is to be expected, as shown in Figure 3A-C, that the generally weaker, high resolution reflections will suffer the greatest proportional disruption due to dynamical scattering. The overall inflation of high resolution intensities reduces the utility of merging statistics such as CC_1/2_ and 〈*I*/*σ*(*I*))〉 in choosing a suitable resolution cut-off. Therefore a relatively conservative criterion of CC_1/2_ ≥ 0.5 was chosen to take this into account. The integrated intensities were then scaled and merged using *AIMLESS* (Evans and Murshudov, 2013).

### Structure solution and refinement

For all datasets, the phases were determined by molecular replacement using Phaser (McCoy et al., 2007). A model of proteinase K (PDB ID: 2ID8) solved by X-ray crystallography was used as a search model with all alternative side chain conformations and ligands removed (Wang et al., 2006). The resultant structures were refined in Phenix using the program phenix.refine with electron scattering factors (Adams et al., 2010). Rounds of refinement were interspersed with manual building using the program Coot (Emsley et al., 2010). Figures of the resultant structures and electrostatic potential maps were generated using Pymol (Schrödinger, LLC., 2015).

### Results and Discussion

#### 1. Sample preparation using cryoFIB milling

Proteinase K was chosen as the test specimen. To reliably generate nanocrystals, seeds (Luft and DeTitta, 1999) were added to batch crystallisation conditions. The larger micrometre-sized crystals used for cryoFIB milling were grown in the same batch conditions by using a larger dilution of the crystal seeds. Both the nanocrystals and the microcrystals (Figure 1A-B) were applied to Quantifoil™ cryoEM grids before vitrification in liquid ethane. The larger proteinase K microcrystals were subject to cryoFIB milling using a Scios™ DualBeam™ instrument following the protocols described in Duyvesteyn *et al*. 2018 (Duyvesteyn et al., 2018) (Figure 1C-E). The resultant lamellae measured approximately 10 μm × 10 μm × 0.2 μm thick. A notable development in the protocol presented here is that the lamellae were sputter coated with metallic Pt after milling to improve sample stability and reduce charging during data collection.

**Figure 1.**
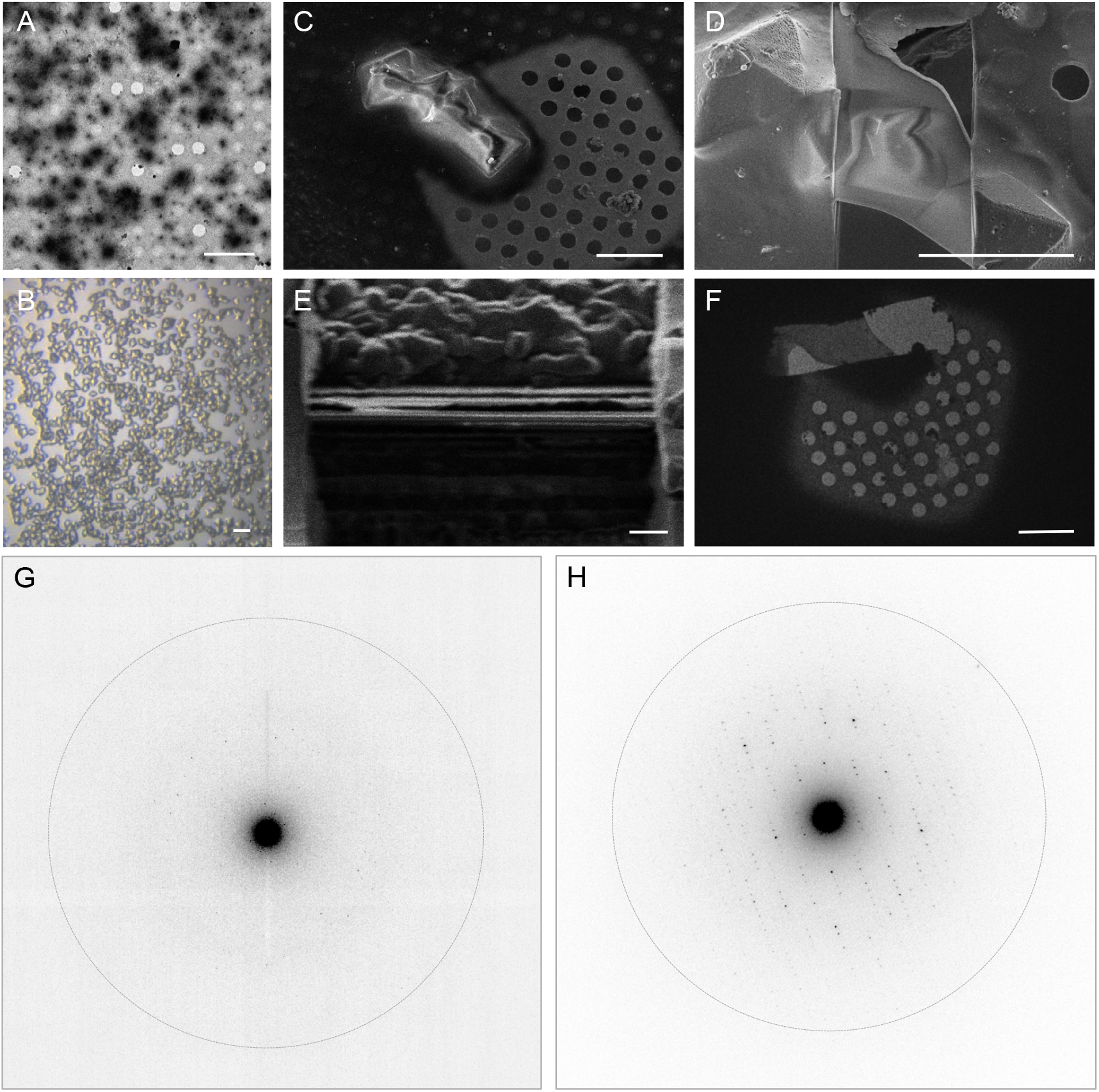
Electron diffraction of proteinase K crystals with and without cryoFIB milling. (A) An electron micrograph of proteinase K nanocrystals. (B) A light micrograph of proteinase K microcrystals. (C-D) Representative SEM images of proteinase K microcrystals before (C) and after (D) cryoFIB milling. (E) An ion beam image of the lamella after the final milling step. (F) A cryoEM image of the resultant proteinase K lamella at low magnification. (G-H) Electron diffraction patterns recorded from proteinase K nanocrystals (G) and from crystal lamella (H). The dotted circle represents the 2.0 Å resolution shell. The scale bars, 10 μm in A, 20 μm in B, 5 μm in C-D, 1 μm in E, and 10 μm in F.

#### 2. MicroED data collection using Ceta-D

Electron diffraction data were collected for both the nanocrystals and the lamellae using the continuous-rotation method (Arndt and Wonacott, 1977). The practical methods described by Shi *et al*. 2016 were used to set up the microscope for data collection with the following modifications: 1) A selected area aperture was not used, instead, a small condenser (C2) aperture of 20 μm was inserted to control the illuminated area which reached the sample. 2) The stage was controlled semi-automatically by a custom program (Supplementary Figure 3) which allowed for continuous-rotation of the sample during exposure to the electron beam without the need for a shutter.

The data were recorded on a Ceta-D detector operating in rolling-shutter mode (Table 1). Using a dose of approximately 0.04 e^−^/Å^2^/s, the Ceta-D was able to successfully measure microED data from nanocrystals to 2.7 Å resolution (Figure 1G), which is consistent with a previous report (Hattne et al., 2019). It is important to note, however, that these nanocrystals were estimated to be at least 500 nm in thickness, and Ceta-D was not sensitive enough to measure diffraction intensities from smaller, thinner crystals using this low-dose setup. In contrast to the nanocrystals, the lamellae measured approximately 200 nm in thickness. Diffraction intensities to 2.4 Å resolution were recorded with the Ceta-D under the same experimental conditions as those used for the nanocrystals (Figure 1H). In addition, we also collected diffraction data from crystal lamellae using a 50 μm C2 condenser aperture, which yielded measurable reflections extending to approximately 2.0 Å (Table 1).

**Table 1.** Data processing, structure solution and refinement statistics for data collected from nanocrystals and from lamella with either a 20 μm or a 50 μm condenser aperture. Numbers in brackets represent the statistics for data in highest resolution shell.

| | Nanocrystals | Lamella (20 $\mu\text{m}$) | Lamella (50 $\mu\text{m}$) |
| --- | --- | --- | --- |
| **Data integration** |  |  |  |
| Space group | $P4_32_12$ | $P4_32_12$ | $P4_32_12$ |
| $a=b, c$ ($\text{\AA}$) | 67.37, 106.78 | 67.33, 106.60 | 67.33, 106.88 |
| $\alpha=\beta=\gamma$ ($^\circ$) | 90.0 | 90.0 | 90.0 |
| No. of datasets | 8 | 4 | 2 |
| No. of crystals | 8 | 2 | 1 |
| Resolution ($\text{\AA}$) | 2.7 | 2.4 | 2.0 |
| $R_{\text{meas}}$ | 0.532 (1.376) | 0.402 (1.323) | 0.332 (1.435) |
| $R_{\text{pim}}$ | 0.140 (0.393) | 0.115 (0.374) | 0.104 (0.450) |
| $\langle I/\sigma(I) \rangle$ | 4.3 (1.8) | 5.2 (2.0) | 5.3 (1.8) |
| Completeness (%) | 88.5 (83.6) | 100.0 (99.7) | 93.0 (92.4) |
| Reflections | 78458 (8010) | 121087 (12723) | 139242 (10323) |
| Unique reflections | 6260 (573) | 10176 (1035) | 15787 (1139) |
| $CC_{1/2}$ | 0.946 (0.498) | 0.987 (0.701) | 0.984 (0.611) |
| **Structure solution** |  |  |  |
| Translation-function Z-score | 41.4 | 50.5 | 59.3 |
| Log likelihood gain score | 2420.355 | 4201.608 | 6312.479 |
| **Refinement** |  |  |  |
| Reflections | 6182 | 10107 | 15758 |
| Reflections used for R-free | 297 | 495 | 792 |
| Resolution range | 56.97-2.70<br>(2.83-2.70) | 56.92-2.40<br>(2.49-2.40) | 56.97-2.00<br>(2.06-2.00) |
| R (%) | 20.92 | 17.75 | 19.40 |
| $R_{\text{free}}$ (%) | 24.82 | 21.59 | 22.68 |
| RMSD bonds | 0.002 | 0.003 | 0.002 |
| RMSD angles | 0.463 | 0.524 | 0.500 |
| $\langle B \rangle$ ($\text{\AA}^2$) | 7.30 | 18.31 | 16.44 |
| Ramachandran plot |  |  |  |
| Favoured (%) | 97.10 | 96.74 | 97.11 |
| Allowed (%) | 2.54 | 3.26 | 2.53 |
| Outliers (%) | 0.36 | 0.00 | 0.36 |
| PDB entry |  |  |  |

#### 3. MicroED data processing using DIALS

The diffraction data were processed using DIALS, which contains implementations specific for electron diffraction (Clabbers et al., 2018). The diffraction images (.ser files) were read directly, without conversion, using the dxtbx library (Parkhurst et al., 2014), while pertinent metadata describing the diffraction experiment were provided separately during data importing. The diffraction images of some data sets displayed negative averaged background levels at higher resolutions (Supplementary Figure 1). This negative bias led to failures during profile fitted integration. The investigation into these failures led to changes to the method used to estimate weights for each pixel used in profile fitting. The new method ensures positive pixel variance estimates, even in the presence of negative-valued pixels. This more robust algorithm also reduces bias in intensity estimates for weak reflections in general and became the default from DIALS version 1.10. Alongside the enhanced profile fitting algorithm, in this specific case we also applied an additive correction to the images to account for the non-physical negative bias. As the magnitude of the bias was less than 1 for all datasets, we added 1 to all pixel values on-the-fly using a dxtbx plugin (Parkhurst et al., 2014), without changes to either the image files or DIALS code.

Four datasets recorded from two lamellae were merged to produce a single dataset that was taken forward to phasing and refinement. Likewise, eight partial datasets from individual nanocrystals were merged to form a complete dataset for further analysis (Table 1). Notably, when using a 50 μm C2 aperture, we were able to produce a complete dataset using just two rotation data collections from a single lamella. For all merged data, molecular replacement was performed with Phaser (McCoy et al., 2007) using proteinase K atomic coordinates determined by X-ray crystallography as a starting model (PDB: 2ID8 (Wang et al., 2006)). Structure refinement, coupled with rounds of manual rebuilding were carried out using Phenix and Coot, respectively (Adams et al., 2010; Emsley et al., 2010). The results from the data processing, structure solution and refinement for the merged datasets are presented in Table 1. Structure solution was successful with the data collected from both the nanocrystals and the crystal lamellae, with example electrostatic potential maps shown in Figure 2. These three structures, derived from nanocrystals, crystal lamellae with 20 μm aperture or crystal lamella with 50 μm aperture, respectively, overlap well (Figure 2A). The resolutions are slightly higher with crystal lamellae (Table 1), with densities subtlely better resolved (Figure 2B-D).

**Figure 2.**
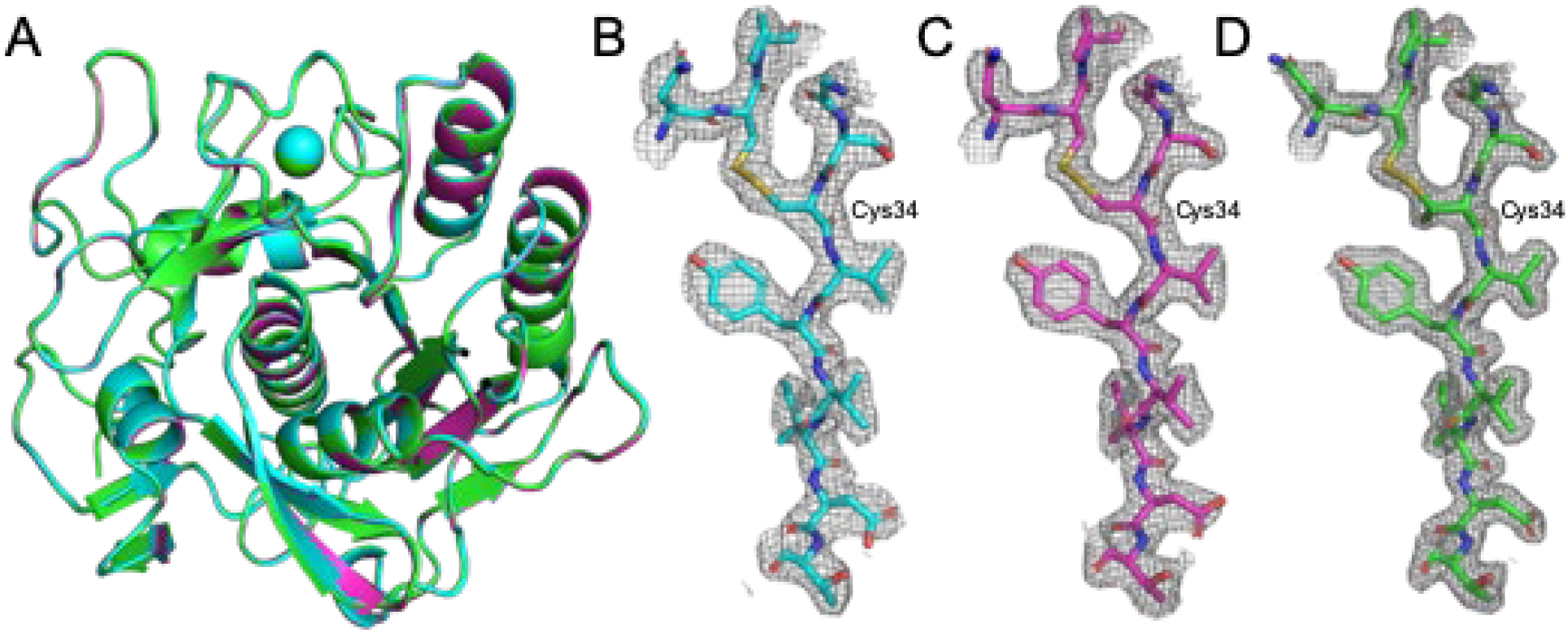
Overall model and example electrostatic potential maps for the proteinase K structures determined using electron diffraction data. (A) The models for the structures determined from nanocrystals (cyan) and the structures determined from lamella with the 20 μm (magenta) and 50 μm (green) condenser apertures are shown aligned by C-alpha residues in cartoon representation with the Ca2+ ion depicted as a sphere. (B-D) A section of the electrostatic potential maps around the disulphide bridge linking residues Cys34 and Cys123 is shown with the 2mFo – Fc maps contoured at 1.0 σ above the mean for the nancrystals (B), 20 μm C2 aperture lamella dataset (C) and the 50 μm C2 aperture lamella dataset (D).

Interestingly, a positive electrostatic potential was present close to the catalytic triad for both the nanocrystal and lamella structure of proteinase K when using 20 μm aperture (Supplementary Figure 2). Existing structures show this to be the binding site for inhibiting Hg atoms (Gourinath et al., 2001; Müller and Saenger, 1993; Saxena et al., 1996), but given that no heavy atoms were present in the crystallisation conditions, these sites were left unmodelled.

#### 4. Comparison of nanocrystals and crystal lamellae

Dynamical scattering in electron diffraction is a concern when crystals are thicker than a few hundred nanometres (Subramanian et al., 2015); still, protein structures have been determined successfully from crystals measuring up to 400 nm thick (Hattne et al., 2015). We analysed the correlation between Fo and Fc for both the nanocrystals, measuring ~500 nm and the crystal lamella, measuring ~200 nm, using the plots described in Clabbers *et al*. (2018) (Figure 3). The nanocrystal data exhibits greater scatter around and deviation from the F_o_=F_c_ line, particularly at low intensity (Figure 3A). This is indicative of dynamical scattering but perhaps also the influence of compound elastic and inelastic scattering events. Interestingly, the average B factor value is considerably lower than would be expected for an equivalent X-ray crystallography structure at this resolution. Indeed, the B factors of several atoms fell to zero during structure refinement. The effect of dynamical scattering on intensities (Voigt-Martin et al., 1997) may emulate the effect of map sharpening and partially explain these low B-factors (Hattne et al., 2015).

**Figure 3.**
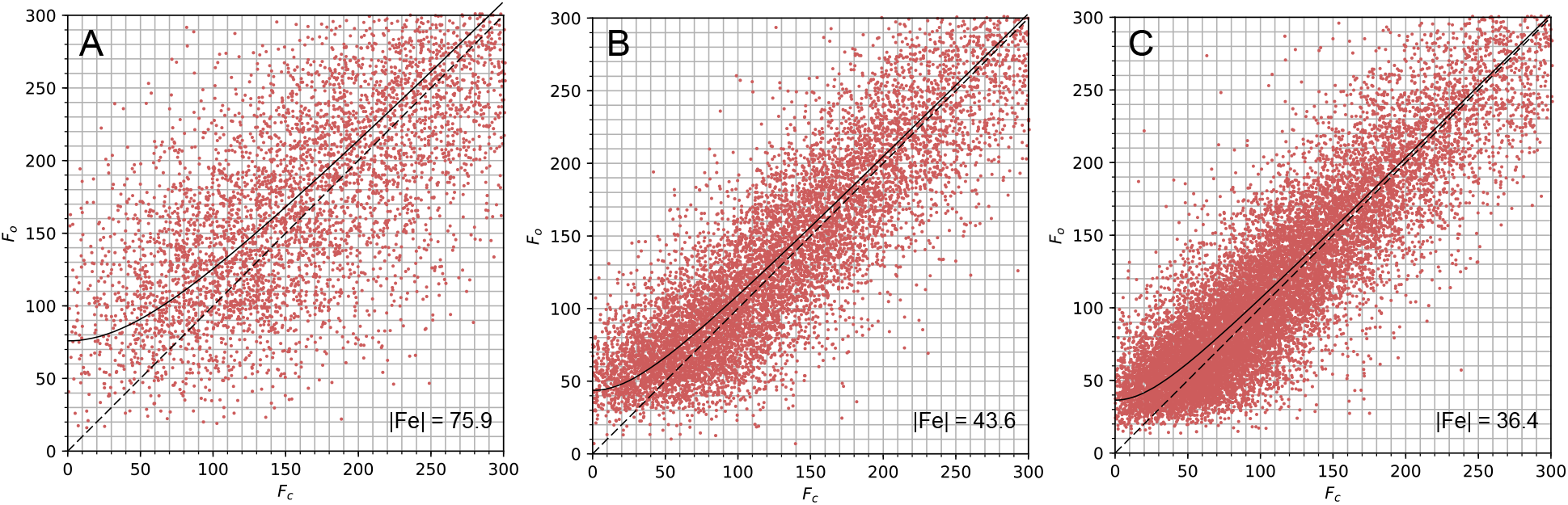
F_o_ vs. F_c_ plots for the proteinase K structures. The Fo vs. Fc plots for the nanocrystals (A) and lamella structures with 20 μm (B) and 50 μm (C) apertures describe the correlation between Fo and Fc for each dataset. The |Fe| value indicates the y-intercept of the curve fitted to these plots and is inset into the bottom right corner of each graph.

The lamella data showed stronger correlation between the Fo and Fc values (Figure 3B and C) suggesting that the data from these samples were less influenced by dynamical scattering. This indicates one of the advantages of using protein crystal lamellae, namely that the sample thickness can be specifically tailored to the requirements of electron diffraction experiments. Higher resolution structures were determined from the lamella data (2.4 Å and 2.0 Å) when compared to the nanocrystal data (2.7 Å). This was likely due to an improvement in signal to noise from the lamella, because the lamella filled the entire illumination area and because neither carbon support nor vitrified mother liquor was present in the exposed area. Furthermore, multiple wedges of data could be acquired from a single crystal lamella as shown here, resulting in better data merging statistics.

Using cryoFIB milled protein crystal lamella opens up the electron diffraction method to all sizes and shapes of crystals, allowing researchers to capitalise on the unique properties of microED, namely electrostatic potential maps and the ability to reveal hydrogen positions, given the data are of sufficient resolution. In addition, microED requires very little sample for protein structure determination compared with macromolecular crystallography (MX) and XFEL methods. Here, we established a streamlined workflow to enable microED for routine protein structure determination. We demonstrated cryoFIB milled lamella give higher resolution data of better quality when compared to nanocrystals under the same imaging conditions. We showed that the low cost Ceta-D camera, generally accessible in a standard cryoEM setup, worked well for microED for well diffracting samples, as reported recently by others (Wang et al., 2019). Future developments to our protocol using a direct electron detectors or a hybrid pixel array detector could potentially improve both data quality and resultant structures, and enable more challenging problems to be tackled.

## Acknowledgements

We thank Dr. Teresa Brosenitsch for critical reading of the manuscript, Dr. Abhay Kotecha and eBIC staff for their technical support. This work was supported by the UK Wellcome Trust Investigator Award 206422/Z/17/Z (PZ) and the UK Biotechnology and Biological Sciences Research Council grant BB/S003339/1 (PZ). We acknowledge Diamond for access and support of the CryoEM facilities at the UK national electron bio-imaging centre (eBIC, proposal NT23169). JMP and DIALS development is supported by Diamond Light Source, CCP4 and Wellcome Award 202933/Z/16/Z (GE).

## Author contributions

G.E. and P.Z. conceived, and with E.V.B. and C.H., designed the experiments. E.V.B., C.H. and J.v.R. prepared crystal samples, C.H, E.V.B. and J.B.G. performed cryoFIB milling of proteinase K crystals, E.V.B., C.H., J.v.R., F.d.H. and B.B. collected electron diffraction data, D.G.W. and J.M.P optimized DIALS software for microED data processing, E.V.B. and D.G.W. analysed data, E.V.B., D.G.W, G.E. and P.Z. wrote the paper with support from other authors.

## Data availability

The datasets generated during and/or analyzed during the current study are available from the corresponding author on reasonable request. Proteinase K structures are deposited at the Protein Data Bank under accession code XXXX for nanocrystals, XXXX for crystal lamellae with 20 μm C2 aperture, and XXXX crystal lamellae with 50 μm C2 aperture.

