## Supplemental figures 1-3 for "A Workflow for Protein Structure Determination from Thin Crystal Lamella by Micro-Electron Diffraction"

for

**This document includes**

Supplementary figures 1-3

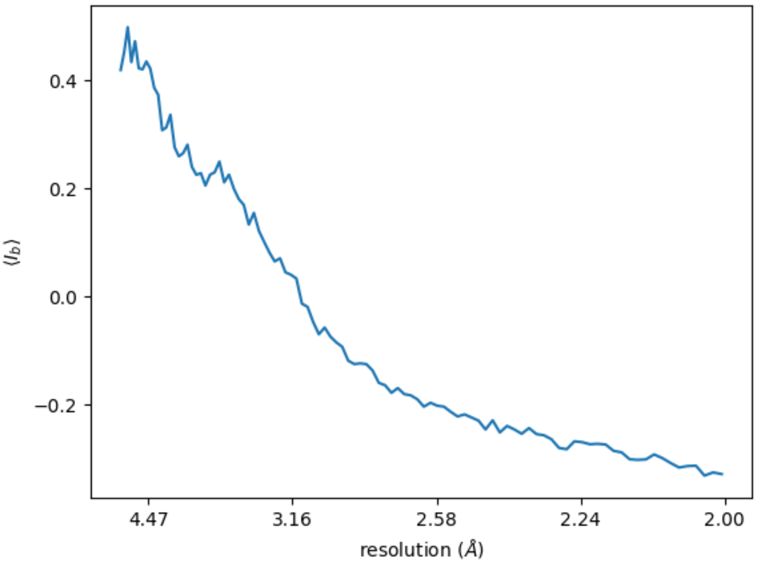


**Supplementary Figure 1 | The average background values from a single frame of a representative nanocrystal dataset.** The average background values were calculated in resolution shells by the program *dials.background* for image 18 of one nanocrystal dataset collected with a 20 µm condenser aperture. This diffraction pattern exhibits the strongest negative bias for that dataset, with average background levels below zero at resolutions beyond about 3.1 Å.

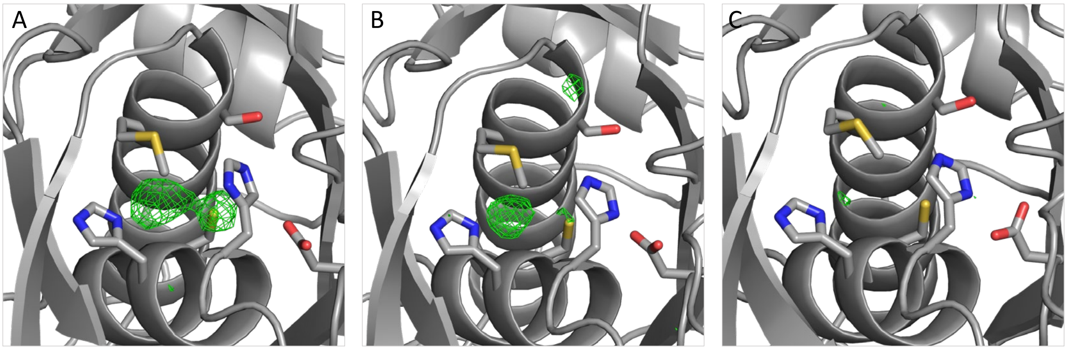


**Supplementary Figure 2 | Positive electrostatic potential near the catalytic triad of proteinase K.** The $m\mathrm{F}_o$-$\mathrm{F}_c$ maps are shown contoured at 3.5 σ above the mean. Positive electrostatic potential can be seen in the nanocrystal maps (A). The lamella maps also show some positive electrostatic potential for the data collected with a 20 µm condenser aperture (B), but not from the data collected with the 50 µm aperture.

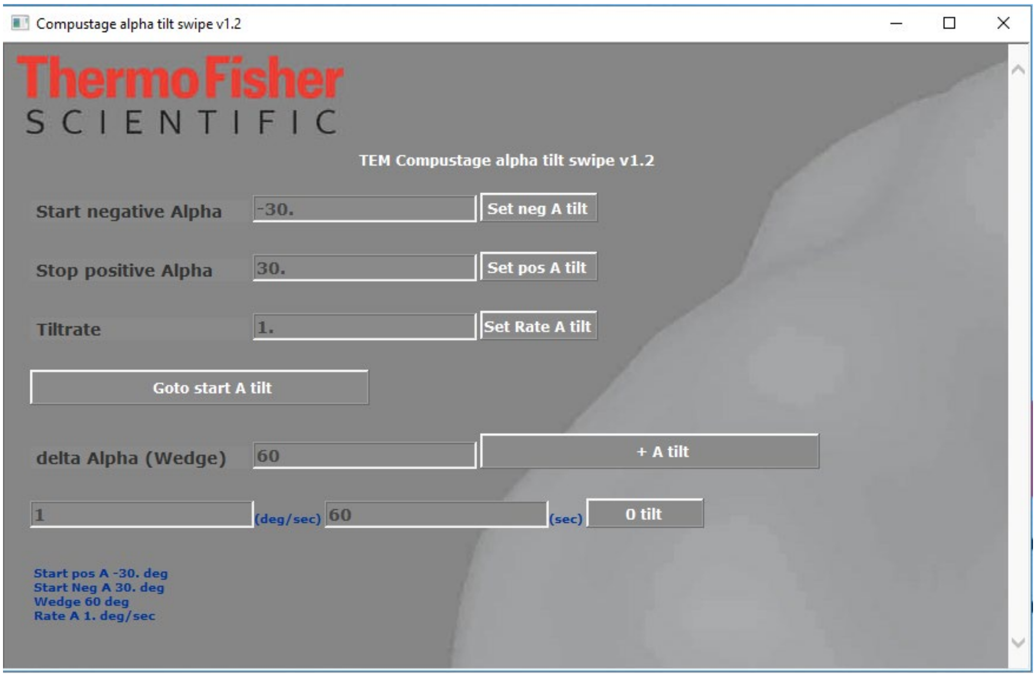


**Supplementary Figure 3 | Continuous rotating electron diffraction data collection script and the user interface.**
